# Metagenomic next-generation sequencing reveals *Miamiensis avidus* (Ciliophora: Scuticociliatida) in the 2017 epizootic of leopard sharks (*Triakis semifasciata*) in San Francisco Bay, California

**DOI:** 10.1101/301556

**Authors:** Hanna Retallack, Mark S. Okihiro, Elliot Britton, Sean Van Sommeran, Joseph L. DeRisi

## Abstract

During March to August of 2017, hundreds of leopard sharks (*Triakis semifasciata*) stranded and died on the shores of San Francisco Bay, California, USA. Similar mass stranding events occurred in 1967 and 2011, yet analysis of these epizootics was incomplete and no etiology was confirmed. Our investigation of the most recent epizootic revealed severe meningoencephalitis in stranded sharks, raising suspicion for infection. On this basis, we pursued a strategy for unbiased pathogen detection using metagenomic next-generation sequencing followed by orthogonal validation and further screening. We show that the ciliated protozoan pathogen, *Miamiensis avidus*, was present in the central nervous system of leopard (*n*=12) and other shark species (*n*=2) that stranded in San Francisco Bay, but absent in leopard sharks caught elsewhere. Whereas this protozoan has previously been implicated in devastating outbreaks in teleost marine fish, our findings represent the first report of a ciliated protozoan infection in wild elasmobranchs. This discovery highlights the benefits of adopting unbiased metagenomic sequencing in the study of wildlife health and disease.

## INTRODUCTION

Mass mortality events among wildlife populations provide insight into ecosystem health and human impact. Rapid die-offs can be caused by physical (e.g. weather), chemical (e.g. toxins), and biological processes (e.g. infectious disease), and may be precipitated by human activity. Further, the underlying cause or the indirect consequences of the die-off itself can pose a risk to human populations. Yet identifying an etiology is often challenging. Conventional approaches to epizootic investigation include environmental assessment, observation of animal behavior, and necropsy with cytology, histology, microbiological and chemical analyses. More recently, for outbreaks of infectious disease, molecular techniques like PCR have enabled rapid and specific pathogen identification. However, PCR tests specific candidates and may be limited by available genetic information. Metagenomic next-generation sequencing (mNGS) provides an unbiased alternative, which has been used successfully in human and animal infections (Dervas et al. 2017; Dill et al. 2017; Langelier et al. 2017; Pfaff et al. 2017; Wilson et al. 2017; Doan et al. 2016; Zylberberg et al. 2016; Wilson et al. 2015; Masuda et al. 2014; Stenglein et al. 2014; Wilson et al. 2014; Stenglein et al. 2012; Kistler et al. 2008). Through the analysis of all nucleic acids in a sample, mNGS can simultaneously test for all known organisms, and can also identify novel pathogens including distantly related species. Furthermore, the cost of NGS technologies continues to decrease, making these methods an increasingly viable option for routine wildlife surveillance and disease investigations.

In the past 50 years, several die-offs of unknown etiology have affected leopard sharks (*Triakis semifasciata*) in San Francisco (SF) Bay, California. In 1967, over 1,000 dead sharks, mainly leopard sharks, were collected in one month in Alameda (Russo and Herald 1968; Russo 2015). More recently, unusual shark deaths were noted in the spring of 2006 (Ota and Media News Staff 2006), and mass mortality again afflicted SF Bay leopard sharks in the spring and early summer of 2011 (Prado 2011; Mertens 2011) involving likely hundreds of leopard sharks though the event was not systematically documented. Moribund sharks were often described as confused and disoriented, with erratic behaviors and swimming patterns.

Scuticociliates are free-living marine protozoa that belong to the subclass Scuticociliatida of the phylum Ciliophora (Gao et al. 2016). As opportunistic pathogens, several species of scuticociliates have been reported to cause disease in diverse marine teleost fish species (Budiño et al. 2011; Munday et al. 1997; Jung, Kitamura, and Song 2005; Moustafa et al. 2010; Iglesias et al. 2001; Whang, Kang, and Lee 2013; Smith et al. 2009; Anderson et al. 2009; Garza et al. 2017; Jung et al. 2011; Turgay and Steinum 2015; Ramos et al. 2007), and recently, in the subclass of cartilaginous fish known as elasmobranchs that includes sharks and rays (Stidworthy et al. 2014; Li et al. 2017). Scuticociliatosis is an economically important problem in the context of commercial marine fish culture (Budiño et al. 2011; Munday et al. 1997; Smith et al. 2009), but has not been observed in wild fish populations to our knowledge.

In this study, we sought to identify a cause for mass mortality of leopard sharks in SF Bay in the spring of 2017. Using mNGS and confirmatory molecular and histologic assays, we identify the scuticociliate *Miamiensis avidus* (*M. avidus*) in the central nervous system of stranded sharks, suggesting that this pathogen could contribute to significant disease in wild elasmobranchs.

## MATERIALS AND METHODS

### Shark stranding surveillance

The majority of shark and ray strandings were reported to the California Department of Fish and Wildlife (CDFW) by members of the public, often via The Marine Mammal Center (Sausalito, CA) and the Pelagic Shark Research Foundation (Santa Cruz, CA). Additional stranding data were provided by East Bay Regional Park District rangers (Oakland, CA), the National Parks Service, and CDFW wardens working in and around San Francisco Bay. Some data were gleaned from iNaturalist (www.iNaturalist.org). Stranding data were also acquired during three brief foot surveys of the Foster City shoreline conducted by CDFW in April, June, and August of 2017. Stranding data included date, location, species, approximate size, condition (live, dead, autolyzed), and abnormal behavior (e.g., swimming in circles). Photos were often submitted, with occasional videos. Stranding data were recorded and sorted on the basis of species and date. Shark and ray strandings were plotted on a map of south San Francisco Bay (**Figure S1**).

### Sample collection

Stranded sharks for necropsy by a CDFW pathologist were chosen based on condition, with preference given to live moribund and fresh dead sharks (non-autolyzed with red gills). Sharks were either necropsied in the field or iced and necropsied at CDFW in Vista, CA within 72 hours of collection. Heads of some leopard sharks were removed and frozen at – 10°C until necropsy. Two additional captive sharks were necropsied: one Pacific angelshark (*Squatina californica*) on display at the Aquarium of the Bay (San Francisco, CA), and one moribund leopard shark on display at the Marine Science Institute (Redwood City, CA). As controls, grossly normal leopard sharks were collected via gill net from Newport Bay in Southern California. An additional great white shark (*Carcharodon carcharias*) and soupfin shark (*Galeorhinus galeus*) were collected from outside SF Bay.

### Necropsy

Sampled sharks were cleaned of external mud and debris via freshwater spray. Species and sex were determined via examination of fins and dentition. Sharks were weighed; total and fork length determined. The dorsum of the head was cleaned and disinfected with multiple passes using disposable Clorox Disinfecting Wipes^®^. When possible, endolymphatic pores were identified. The endolymphatic fossa (oval concave depression in the chondrocranium) was located by digital palpation. Using sterilized instruments, a 3×5cm incision was made centered on the endolymphatic fossa and pores. Subcutaneous tissues, overlying the fossa, were sampled with a sterile cotton swab for microbiologic assessment. Subcutaneous fluid was aspirated with a sterile 1mL pipette for cytological assessment. The skin sample containing the endolymphatic pores and ducts was fixed in 10% formalin. The calvarium, including the endolymphatic fossa, was removed with a sterile scalpel and new blade, then fixed in formalin. Removal of the calvarium exposed both inner ears and the cerebellum. Cerebrospinal fluid (CSF) overlying the cerebellum was sampled with a sterile cotton swap. Two 1mL CSF samples were taken by sterile pipette and frozen at –10°C in cryovials. A third CSF sample was taken for cytological assessment. Perilymph from one inner ear was sampled with a sterile cotton swab. A second perilymph sample was taken for cytological assessment. The brain and olfactory lamellae were exposed via sharp dissection. The meninges, CSF, brain, inner ears, and olfactory lamellae were examined for evidence of inflammation and hemorrhage. Brains were separated from the chondrocranium via inversion of the skull and severing cranial nerves. Olfactory lamellae and associated olfactory bulbs were removed via sharp dissection. The brain and olfactory lamellae were fixed in 10% formalin. In some sharks, one otic capsule was also taken and fixed in formalin. Gills, heart, kidneys, and abdominal organs were also examined at necropsy. Selected organs were sampled and fixed in formalin from some sharks.

### Cytology

Samples of subcutaneous fluid surrounding the endolymphatic ducts, inner ear perilymph, and CSF were examined on glass slides under darkfield light microscopy at 200 and 400X using a binocular microscope. Red blood cells, inflammatory cells, and microbial pathogens were identified.

### Histology

Histology samples were immersion fixed in 10% formalin for 2 weeks to 3 months and then routinely paraffin processed. Paraffin blocks were sectioned at 5-7μm and sections stained with hematoxylin and eosin (HE), then examined with light microscopy for inflammation, necrosis, and pathogen presence.

### Microbiology

Samples of subcutaneous fluid surrounding the endolymphatic ducts, inner ear perilymph, and CSF were plated onto blood agar (for bacterial pathogens) and Sabouraud-Dextrose agar (for fungal pathogens). Cultures were incubated aerobically at room temperature (15-20°C) for 4 weeks and checked daily for growth. Selected isolates were sent to the University of Florida (Gainesville, FL) for biochemical and PCR identification.

### Nucleic acid extraction

For RNA, 250μL of CSF was placed in TRI-Reagent (Zymo Research) and homogenized with 2.8mm ceramic beads (Omni) on a TissueLyser II (Qiagen) at 15Hz for two 30sec pulses, separated by 1min on ice. Total RNA was then extracted using the Direct-Zol RNA MicroPrep Kit with DNase treatment (Zymo Research), eluted in 12μL and stored at –80°C until use. For DNA, 250μL of CSF was placed in 750μL Lysis Solution of the Fungal/Bacterial DNA kit (Zymo Research) and homogenized as above with a single 2min homogenization pulse. Total DNA was then extracted using the Fungal/Bacterial DNA kit (Zymo Research), eluted in 25μL, and stored at –80°C until use.

### Sequencing library preparation

RNA samples were processed using 5μL total RNA as input into the NEBNext Ultra II RNA Library Prep Kit for Illumina (New England Biolabs). Samples were sequenced on an Illumina MiSeq instrument using 150 nucleotide (nt) paired-end sequencing. A “no-template control” (nucleic acid-free water) was included in each batch of nucleic acid extractions and library preparation. Raw sequencing reads were deposited at the National Center for Biotechnology Information (NCBI) Sequence Read Archive (SRA) under BioProject PRJNA438541, SRA accession SRP136047.

### PCR/Sanger sequencing

Primers used to amplify ciliate and shark genomic sequences are listed in **Table S3**. PCRs were performed in a final volume of 50μL containing 1X Phusion HF buffer (New England Biolabs), 0.2mM dNTPs, 0.5μM each primer, 1U of Phusion DNA polymerase (New England Biolabs) and template DNA (1μL of extracted DNA for FISH5.8SF/FISH28SR reactions, 2 μL for OX09-26/OX09-27, or 5μL for Cil3/Cil4 reactions). Samples were denatured for 1min at 98°C, then cycled with 10sec denaturation at 98°C, 15sec annealing, and 30sec extension at 72°C, followed by a final 5min extension at 72°C. For primer pairs (i) FISH5.8SF/FISH28SR (Pank et al. 2001), (ii) OX09-26/OX09-27 (Whang, Kang, and Lee 2013), and (iii) Cil3/Cil4 (Jung, Kitamura, and Song 2005), annealing temperatures were (i) 72°C, (ii) 55°C, and (iii) 52°C, extension times were (i) 2min, (ii) 30sec, and (iii) 30sec, and (i) 30, (ii) 35, or (iii) 35 cycles were performed. Nested PCR was performed with 1/150^th^ of purified OX09-26/OX09-27 reaction products (purified with DNA Clean & Concentrator-5 columns (Zymo Research) to remove primers), in 20μL reactions scaled from the conditions above, using ciliate species-specific primers OX09-142 through OX09-149 (Whang, Kang, and Lee 2013), with annealing temperature of 60°C and extension time of 30sec, for 30 cycles. Amplified products were visualized on a 1.5% (Cil3/4 and ciliate species-specific reactions) or 1% (FISH5.8SF/FISH28SR and OX09-26/OX09-27 reactions) agarose gel stained with GelRed (BakerBiotium) under UV light. Purified PCR products from OX09-26/OX09-27 and Cil3/Cil4 reactions were sequenced via the Sanger method by QuintaraBio (Albany, CA).

### Bioinformatics

#### Metagenomic Next-Generation Sequencing (mNGS)

Next-generation sequencing data were analyzed using a computational pipeline originally developed by the DeRisi Laboratory to identify potential pathogens in human samples (Michael R. Wilson et al. 2014). Briefly, host sequences were identified using publicly available shark genome/transcriptomes, and remaining non-host sequences were compared to the NCBI nucleotide and protein databases. Potential pathogens were identified based on a minimum read abundance, likelihood of pathogenicity, and absence in negative control samples. For species determination, reads mapping to the ciliate 18S small subunit (SSU) and 28S large subunit (LSU) of the nuclear ribosomal RNA locus (rDNA) were assembled and compared to the NCBI database using BLASTn (Altschul et al. 1990). See Supplemental Methods for details.

#### Phylogenetic sequence analysis

New sequences in this study include the partial sequences of the mitochondrial (mt) cytochrome c oxidase I (*cox1*) gene of ciliates from seven shark samples (by Sanger, labeled by Fish ID); the SSU rDNA sequence (by Sanger – deposited sequence is 100% identical to sequences from five shark samples, and additional consensus contigs from NGS); and the LSU rDNA (consensus contigs from NGS), deposited in GenBank (Accession numbers MH078243-MH078249, MH062876, MH064355). For comparison, *cox1*, SSU, and LSU sequences of representative philasterid species (**Tables S4-S6**) were downloaded from GenBank and aligned with the sequences identified in this study using Geneious (Biomatters Ltd, v9.1.8). Trimming ends to the shortest common region gave a total of 643, 958, and 1,799 sites for *cox1*, SSU, and LSU respectively, which were used to construct NJ phylogenetic trees in Geneious (Biomatters Ltd, v9.1.8) using the Tamura-Nei distance method with *Tetrahymena pyriformis* as an outgroup. The confidence estimates in the NJ phylogenetic trees were determined by 1000 bootstrap re-samplings.

## RESULTS

### Stranded leopard sharks display inflammatory meningoencephalitis

Beginning in March 2017, members of the public reported sharks swimming with unusual behaviors and stranding on beaches along the SF Bay shoreline, with the majority of strandings occurring in the Foster City area (**Figure S1**). Leopard sharks were observed swimming unusually close to shore, appearing uncoordinated and disoriented, suggestive of an inner ear or central nervous system issue. At the height of the epizootic in April and May, 20-30 dead leopard sharks were being found daily along the shoreline in Foster City shoreline, and we estimated that over 1,000 leopard sharks died between March and August 2017 in SF Bay.

Necropsies were performed on 11 fresh dead or live moribund leopard sharks, and on the heads of five frozen sharks. Gross and cytological lesions were consistent with meningoencephalitis, and characterized by hemorrhage, cloudy CSF, and thickened meninges (Figure 1, Table 1, **Table S1**). Lesions were especially prominent in the olfactory bulbs and lobes. Olfactory lamellae, adjacent to olfactory bulbs, were often markedly hemorrhagic and inflamed. There was no gross evidence of inflammation in the subcutaneous tissues surrounding the endolymphatic ducts or inner ears, which are target organs for a common bacterial pathogen of sharks (*Carnobacterium maltaromaticum*) (Schaffer et al. 2013). No lesions were observed in gills, heart, or abdominal organs. Cytologic exam of CSF revealed dense mixed inflammation (mononuclear inflammatory cells and polymorphonuclear cells). No pathogens were observed. Conventional microbiology was uninformative: blood and Sabouraud-Dextrose agar cultures of CSF, inner ear perilymph, and subcutaneous tissues and surrounding endolymphatic ducts yielded no growth or fungal and bacterial contaminants associated with field sampling or post-mortem colonization of tissues.

**Figure 1:**
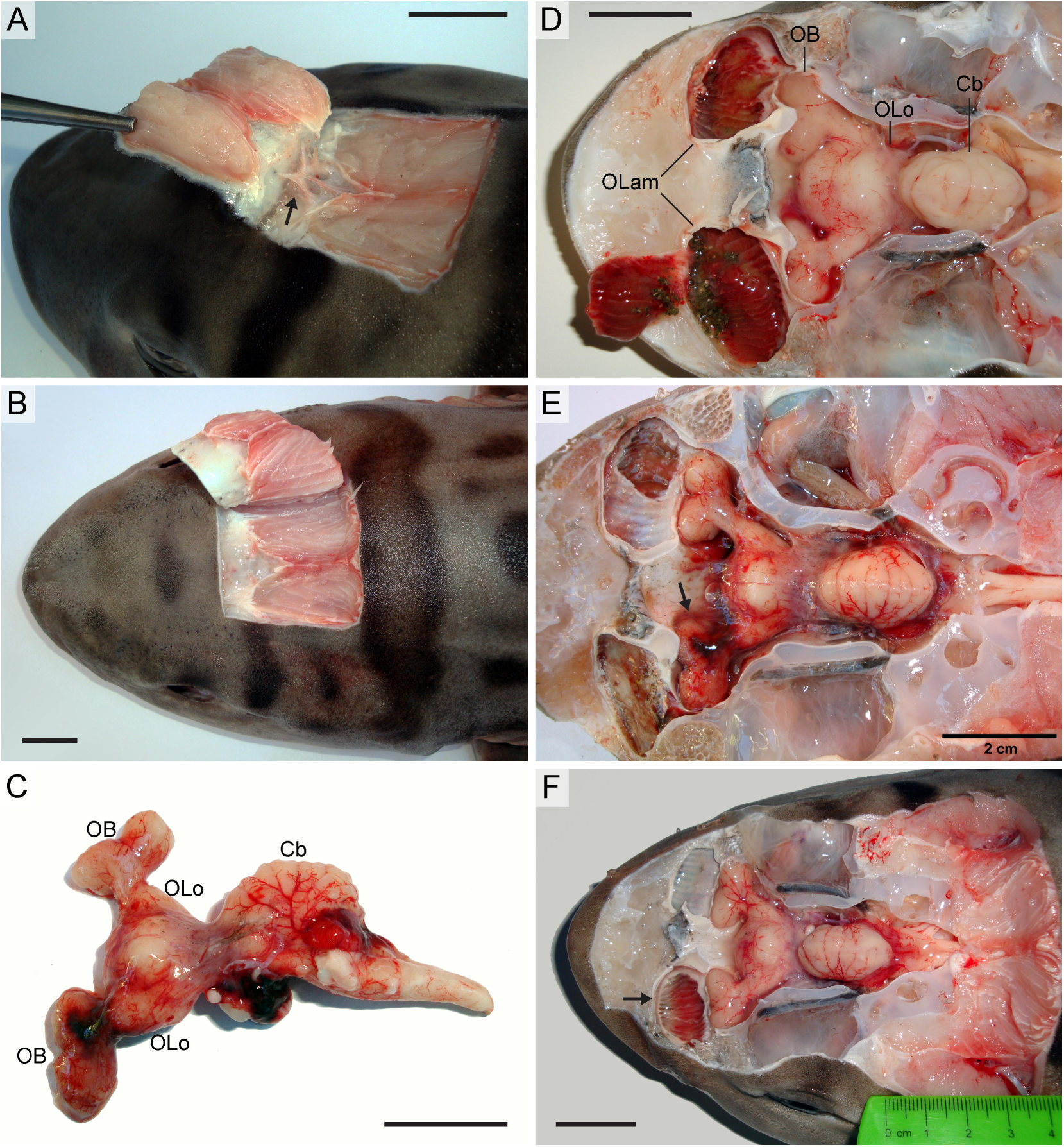
Gross histology and lesions in brains of stranded leopard sharks. **A-B)** Dissection from the superior aspect of the head exposing the endolymphatic ducts (arrow). **C)** Brain with hemorrhagic lesions removed from cranial cavity of (B) and depicted in situ in (E). **D-F)** Dissection of cranial vault revealing superior surface of brain, hemorrhagic lesions and congested olfactory lamellae (arrows). (OB) olfactory bulb; (OLo) olfactory lobe; (Cb) cerebellum; (OLam) olfactory lamellae. Scale bars, 2cm.

**Table 1.**
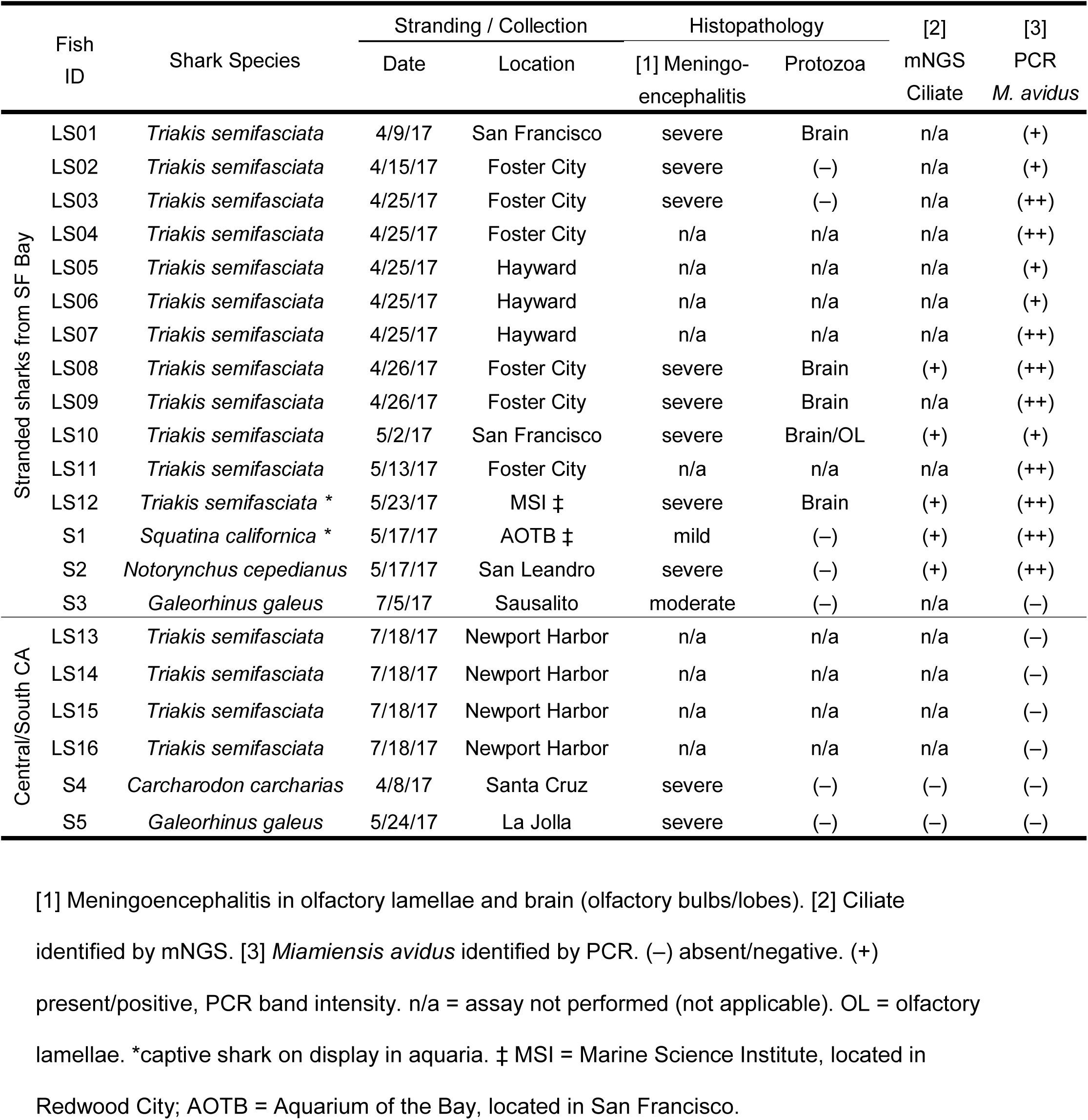
Summary of results. Sharks grouped by involvement in epidemic (stranded sharks from SF Bay vs. controls from Central/Southern CA).

Guided by this picture of meningoencephalitis, samples of CSF were taken for molecular analysis from 15 stranded or ill-appearing sharks from SF Bay including 11 leopard sharks (*Triakis semifasciata)*, one sevengill shark (*Notorynchus cepedianus*), and one soupfin shark (*Galeorhinus galeus)*, as well as one captive leopard shark from the Marine Science Institute (Redwood City) and one captive Pacific angelshark (*Squatina californica*) from the Aquarium of the Bay (San Francisco) (Table 1, **Table S1**). Control CSF samples were collected from four grossly normal leopard sharks captured by gill net from Newport Bay in Southern California, and from two sharks with meningoencephalitis deceased in Southern or Central California: one great white shark (*Carcharodon carcharias*) and one soupfin shark.

### mNGS reveals scuticociliate sequences in cerebrospinal fluid (CSF) of stranded sharks

To identify potential pathogens associated with leopard shark mortality, we performed mNGS on CSF samples from five sharks exposed to SF Bay water and two sharks from elsewhere on the California coast (Table 1). Reads aligning to species in the Ciliophora phylum (taxonomy ID 5878) were identified in all five SF Bay sharks, but were absent from the no-template control and non-SF Bay sharks (**Table S2**). No other credible pathogens were identified. Sixty-four percent (64%) of identified Ciliophora reads aligned to the ciliate rDNA locus. High-confidence contigs were assembled for the partial SSU (943nt with a gap of 310nt) and LSU (1916nt with gaps of 39nt and 56nt) rDNA, with coverage between 4 and 2545 unique reads, and nucleotide identity >99% except at 2 loci: nt 927 of SSU (88% T, 10% G) and nt 1437 of LSU (86% T, 9% C) (numbering per deposited sequences). The SSU contig aligned with greater confidence to *Miamiensis avidus* (*M. avidus*) than other scuticociliate species by BLASTn.

### Histopathology and molecular characterization supports *M. avidus* infection

To confirm mNGS results with an orthogonal molecular approach, we amplified a variable region of the ciliate *cox1* gene from DNA extracted from CSF of these sharks. Amplification was detected in a nested PCR for *M. avidus* in five of five stranded sharks that were positive for *M. avidus* by mNGS (Figure 2A, **Figure S2**), and no amplification was detected in two of two sharks negative by mNGS or in the no-template control (**Figure S2**). The sequence of the *cox1* amplicons (via the Sanger method) was most similar to other *M. avidus* sequences (Figure 2B).

**Figure 2:**
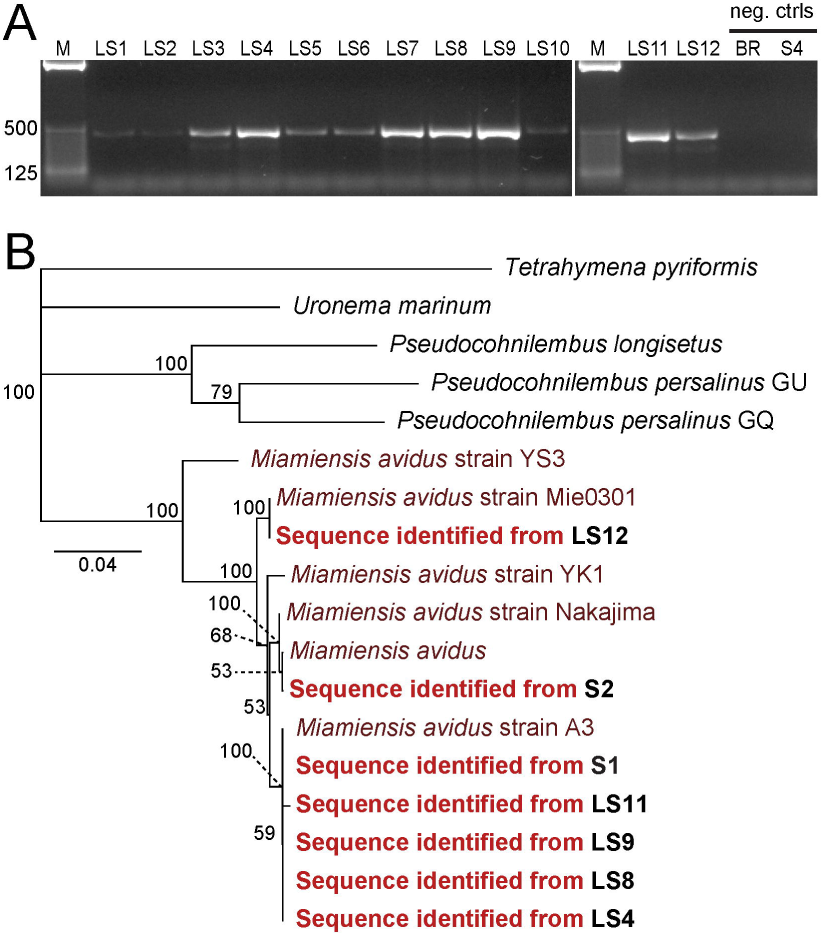
Identification of *M. avidus* in CSF from leopard sharks in SF Bay. **A)** DNA samples from SF Bay leopard sharks (LS1-12) and negative control animals (bat ray (BR) and S4) were tested by nested PCR using primers specific to the *cox1* gene of *M. avidus* (expected size 422bp). M: 25bp ladder. B) Neighbor-joining phylogenetic tree constructed from mt *cox1* nucleotide sequences. *Tetrahymena pyriformis* served as the outgroup. New sequences in this study are in bold, labeled according to Fish ID (see Table 1). Nodes are labeled with bootstrap values based on 1,000 resamplings. See **Table S4** for accession numbers of reference sequences. Scale bar, nucleotide substitutions per site.

In addition, brain and nasal tissues from nine affected sharks were examined for histopathologic evidence of ciliated protozoa. Severe inflammatory and necrotizing lesions were observed in the olfactory bulbs and lobes of the brains, in addition to severe inflammation in the olfactory lamellae (Table 1, Figure 3). Ciliated protozoan parasites, morphologically consistent with *M. avidus*, were observed in the brains of five affected sharks, and in olfactory lamellae of one shark (Table 1, Figure 3).

**Figure 3:**
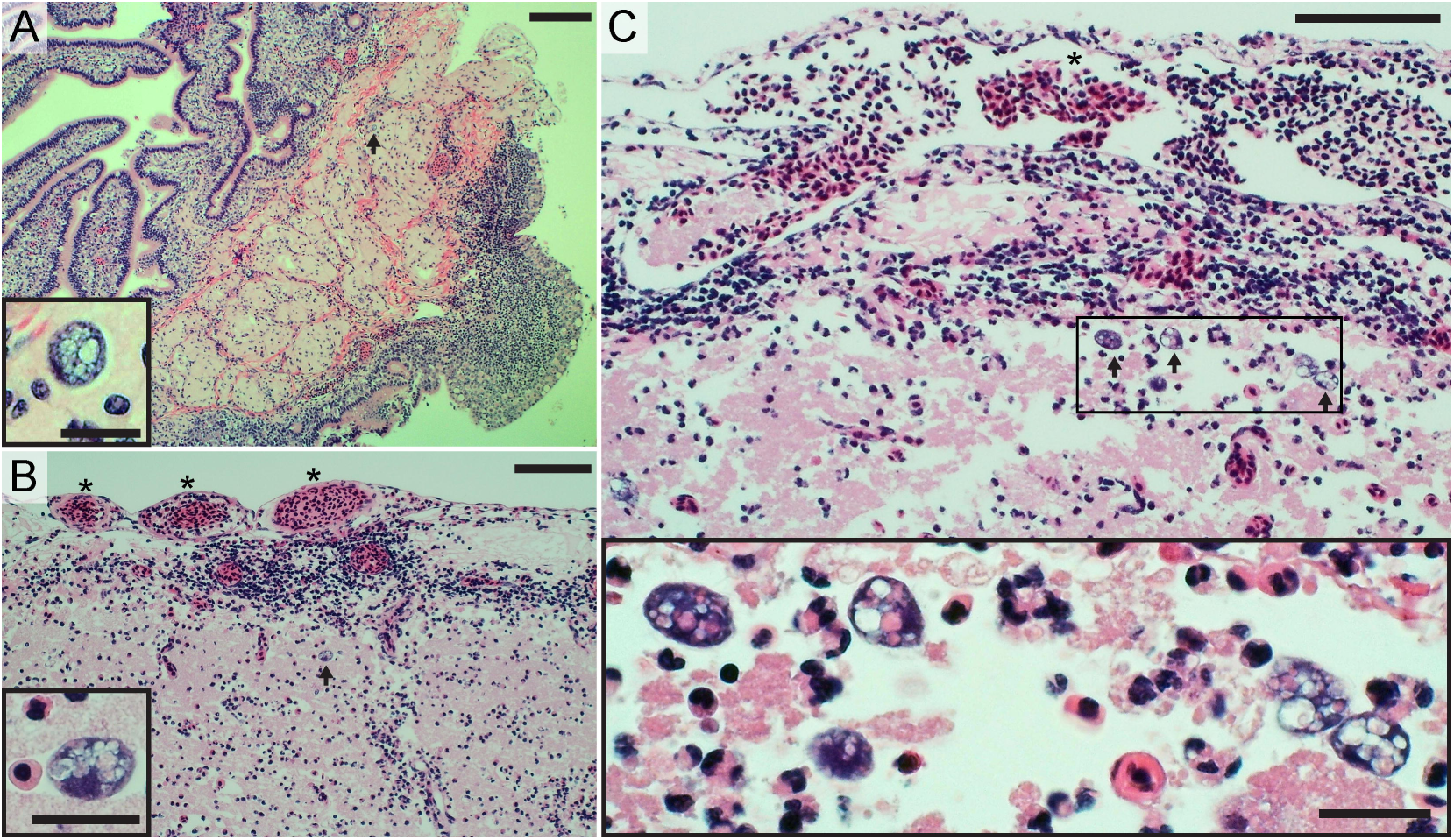
Protozoa identified in brain tissues of stranded leopard sharks. Representative hematoxylin and eosin stained histology sections from stranded leopard sharks. **A)** Olfactory lamellae (top left) and filament (center), with submucosal inflammation (bottom right) and scattered protozoa (arrow, magnified in inset). Scale bars, 200μm and 25μm (inset). **B-C)** Olfactory bulb of the brain with congested vessels in overlying meninges (asterisks), inflammatory infiltrate, and protozoa (arrows, magnified in insets). Scale bars, 100μm and 25μm (insets).

### Presence of *M. avidus* is strongly associated with the SF Bay epizootic

With evidence for *M. avidus* as a candidate pathogen, we screened nine additional leopard sharks that stranded in SF Bay in spring of 2017, compared to four grossly normal leopard sharks that were caught in a gill net in Southern California. A soupfin shark (*Galeorhinus galeus*) that stranded in SF Bay in July 2017 was also tested. PCR was targeted to the ciliate SSU and mt *cox1* regions, initially using universal ciliate primers (Jung, Kitamura, and Song 2005; Whang, Kang, and Lee 2013), followed by Sanger sequencing and/or species-specific nested PCR for the *cox1* gene to test for the related pathogenic ciliate species: *Uronema marinum, Pseudocohnilembus longsietus*, and *Pseudocohnilembus persalinus* (Whang, Kang, and Lee 2013). For *cox1*, amplification specific to *M. avidus* was detected in nine of nine SF Bay leopard sharks (Figure 2A, **Figure S3**), and was not detected in the Southern California leopard sharks (**Figure S4**) or the SF Bay soupfin shark (**Figure S2**). Amplicon sequencing revealed 99.1% pairwise identity. The *cox1* sequences clustered together with reference *M. avidus* sequences on a neighbor-joining (NJ) tree (**Figure 2B**). For the SSU, an amplicon of the expected size was detected in three of 12 SF Bay leopard sharks and in the two SF Bay non-leopard sharks that were positive by mNGS, and was absent from four of four leopard sharks and two of two non-leopard sharks from Southern California (Figure S5). The sequences of the ciliate SSU amplicon from five sharks (LS3, LS4, LS11, S1, and S2) were 100% identical, were concordant with the regions of overlap from mNGS, and clustered together with reference *M. avidus* sequences on a NJ tree (**Figure S6**).

## DISCUSSION

In this study, we describe an epizootic of wild leopard sharks characterized by stranding behavior and meningoencephalitis, and provide strong molecular and histological evidence that implicates the ciliated protozoan, *M. avidus*, as the candidate pathogen associated with the SF Bay mass mortality event. We identified *M. avidus* through an unbiased, NGS-based approach, which we and others have previously used in a wide range of human and non-human infectious disease investigations (Dervas et al. 2017; Dill et al. 2017; Langelier et al. 2017; Pfaff et al. 2017; Wilson et al. 2017; Doan et al. 2016; Zylberberg et al. 2016; Wilson et al. 2015; Masuda et al. 2014; Stenglein et al. 2014; Wilson et al. 2014; Stenglein et al. 2012).

Unlike our previous investigations, we encountered a technical challenge in analyzing mNGS data from a host species lacking a complete genome. To our knowledge, the leopard shark genome has not been sequenced, and thus we utilized available sequence from related species. Despite being unable to identify all host sequences using our proxy-metagenome host sequences, we were still able to identify a plausible pathogen embedded in a large amount of unrelated, unidentified host sequence. Future contributions of shotgun sequencing data including this study will improve our ability to identify sequences of unusual hosts, such as sharks, thereby improving our ability to detect novel pathogens. The need for high-quality samples to preserve RNA and minimize contamination is especially clear in the context of fish mortality, where samples may originate in the field where death is followed by rapid colonization and environmental exposure. When high-quality samples are unavailable, PCR offers a good alternative for pathogen identification. Among the highly-variable regions flanked by conserved sequences, such as the commonly used ribosomal RNA (SSU and LSU) and *cox1* genes, we found that *cox1* was similar to SSU and better than LSU in discriminating between *M. avidus* and related pathogenic scuticociliates. This finding is consistent with reports of higher intraspecific variation of the *cox1* gene (Jung et al. 2011; Budiño et al. 2011). Nonetheless, using multiple genes for molecular phenotyping can add confidence, as there remain discrepancies in the field about highly-similar taxa such as *M. avidus* and *Philasterides dicentrarchi* (Jung et al. 2011; De Felipe et al. 2017), and there are likely sub-species divisions yet to be realized (Gao, Katz, and Song 2012).

We observed *M. avidus* only in sharks exposed to SF Bay water, including two wild-caught animals in captivity. The associated phenotype is consistent with other reports of ciliate infection of elasmobranchs, notable for necrotizing meningoencephalitis (Stidworthy et al. 2014; Li et al. 2017). Pathogenesis in leopard sharks likely involves a nasal route, with initial protozoal invasion of olfactory lamellae, followed by extension into the olfactory bulbs and lobes of the brain. Massive inflammation and severe encephalomalacia, associated with *M. avidus* infection, could account for the disorientation and abnormal behavior of sharks prior to stranding. Further support implicating *M. avidus* in these repeated die-offs comes from the necropsy of a single leopard shark that stranded in the 2011 SF Bay epizootic, which showed extensive inflammation and an abundance of unicellular ciliated protozoa throughout the brain (communication from S. Kubiski, veterinary pathology resident, University of California, Davis, Pathology #11N1368 Final Report, July 1, 2011). We found no evidence of other pathogens that have been reported in elasmobranchs near the Pacific coast of North America (Schaffer et al. 2013; Méndez and Galván-Magaña 2016; Benz, Borucinska, and Greenwald 2009).

Nonetheless, the data do not exclude the possibility that *M. avidus* was not the primary or sole driver of disease and mortality. Factors such as water temperature, salinity, toxins, or other pathogens could increase susceptibility to an opportunistic infection by *M. avidus* (Hobbs, Cook, and Crain 2015; Carlisle and Starr 2009; Hopkins and Cech 2003). In SF Bay, leopard sharks may be especially vulnerable each spring when they aggregate in large numbers in the warm shallow waters of bays and estuaries, with greater exposure to runoff that may contain toxins or decreased salinity (Nosal et al. 2013; Hight and Lowe 2007). Notably, heavy seasonal rainfall with runoff into the bay preceded each of the spring epizootics in 2006, 2011, and 2017 (**Figure S7**) (NOAA National Centers for Environmental Information 2018). Although the specific exposures are unknown, it would be prudent to monitor marine fish hatcheries for scuticociliates, as outbreaks are a known issue in such settings and may pose a risk to wild marine fish stocks. Further studies are needed to clarify susceptibility factors and exposures, especially in the context of major urban centers where planned human development could prevent or mitigate negative impacts of human activity on wild marine fish.

While this epizootic was easily noticed in SF Bay where large numbers of sharks washed up on beaches regularly surveyed by the public and wildlife researchers, scuticociliatosis may also affect elasmobranchs or other marine animals elsewhere along the California coast. Technologies to remotely monitor inaccessible areas and connect casual observers with dedicated researchers could enable better surveillance and minimize unnecessary human intrusion into wild habitats. Future investigations of mass mortality events should include *M. avidus* as a potential pathogen. We anticipate that the episode of scuticociliatosis in wild elasmobranchs described here is not an isolated event. As similar epizootics are uncovered through seasonal monitoring, future research is needed to describe the host-pathogen relationship and potential implications for nearby human populations. Although the only known ciliate parasite of humans, *Balantidium coli*, is far distantly related to scuticociliates (Schuster and Ramirez-Avila 2008), and there are no documented infections of mammals by scuticociliates, sport fishing and consumption of leopard sharks is common in SF Bay and the consequences of *M. avidus* ingestion are unknown. Finally, this study demonstrates the ability of mNGS to rapidly identify potential pathogens in an unbiased manner. Surveillance and disease investigations in wildlife populations will likely benefit from the incorporation of mNGS-based techniques.

## ACKNOWLEDGEMENTS

We would like to acknowledge the Marine Science Institute (Redwood City, CA), the Aquarium of the Bay (San Francisco, CA), The Marine Mammal Center (Sausalito, CA), East Bay Regional Park District (Oakland, CA), the National Parks Service, Jennifer Kampe, and Paige Coluccio for their assistance in this work. We would also like to acknowledge Eric Chow and Derek Bogdanoff at the Center for Advanced Technology (UCSF) for assistance with sequencing. This work was supported by the Chan Zuckerberg Biohub (JD), UCSF Medical Scientist Training Program (HR), and the State of California (MO).

